# Genetic encoding and expression of RNA origami cytoskeletons in synthetic cells

**DOI:** 10.1101/2024.06.12.598448

**Authors:** Mai P. Tran, Taniya Chakraborty, Erik Poppleton, Luca Monari, Franziska Giessler, Kerstin Göpfrich

## Abstract

The central dogma at the core of molecular biology states that information flows from DNA to RNA and then to protein. Our research seeks to introduce a conceptually novel approach towards synthetic life by leveraging RNA origami, as an alternative to proteins, requiring only a single copying step between genetic information and function. Here, we report the genetic encoding and expression of an RNA origami cytoskeleton-mimic within giant unilamellar lipid vesicles (GUVs). We design the first RNA origami tiles which fold co-transcriptionally from a DNA template and self-assemble into higher-order 3D RNA origami nanotubes at constant 37 *^◦^*C in GUVs, where they reach several micrometers in length. Unlike pre-formed and encapsulated DNA cytoskeletons, these GUVs produce their own molecular hardware in an out-of-equilibrium process fuelled by nucleotide feeding. To establish genotype-phenotype correlations, we investigate how sequence mutations govern the contour and persistence length of the RNA origami nanotubes with experiments and coarse-grained molecular-dynamics simulations, realizing a phenotypic transition to closed rings. Finally, we achieve RNA origami cortex formation and GUV deformation without chemical functionalization by introducing RNA aptamers into the tile design.Altogether, this work pioneers the expression of RNA origami-based hardware in vesicles as a new approach towards active, evolvable and RNA-based synthetic cells.

## 1 Introduction

Bottom-up synthetic biology has gathered an interdisciplinary scientific community behind the visionary goal of engineering a cell—the minimal unit of life—from nonliving molecular components [1]. Central to this effort is the imperative for synthetic cells to anabolically construct their own hardware from an intrinsic genome and environmental nutrients, which would support evolutionary adaptation through natural selection. In order to achieve this, each synthetic cell must be equipped with some form of an information-bearing molecule which correlates with cellular function. The most obvious approach is to leverage the information-function relationship that biology already developed, namely the “Central Dogma of Molecular Biology”, which states that information in biological systems flows from DNA (information) to RNA (amplification and regulation) and finally to protein (function) [2] (Fig. 1a). Entrenched in nature’s evolutionary processes, this pathway remains the only known, fully-functional chassis for life and would therefore appear to be a good starting point for the bottom-up assembly of a synthetic cell [3].

Protein production according to the central dogma with cell-free *in vitro* transcription-translation (IVTT) systems has already been demonstrated inside of giant unilamellar vesicles (GUVs) [4–7]. However, protein production in bottom-up assembled synthetic cells comes with several challenges such as low encapsulation efficiency of IVTT systems, limited post-translation modification and glycosylation, cross-talk between components, and life spans limited by ATP regeneration systems or saturated translation machinery [8]. Moreover, we have to acknowledge that a synthetic genome that follows the conventional central dogma would require over 150 genes for the assembly of the transcription-translation machinery alone [3, 9]. The realization of a minimal self-replicating IVTT system has therefore become an active field of investigation on its own [10].

The challenges related to the construction of synthetic cells according to the central dogma encourages a search for alternate solutions. At the onset of life, simpler processes must have sustained self-replication and evolution [11]. One potential solution is to “short circuit” the Central Dogma and create a system with only a single step between the genotype and phenotype molecules. DNA nanotechnology, fuelled by the development of the DNA origami technique [12], has facilitated the creation of structural and, to some extent, functional analogs of cellular proteins that have been reconstituted inside of lipid vesicles. These represent initial steps towards a non-protein-based machinery for synthetic cells [13]. Examples include DNA-based transmembrane pores for substrate exchange [14, 15] or signalling [16], mimics of SNARE proteins for vesicle fusion [17, 18] or scaffolding proteins to generate membrane curvature [19, 20], cell-cell linkers for vesicle populations [21, 22], and condensates to sequester reactions [23] or mimic DNA segregation [24]. Particularly relevant to this work, DNA nanotubes as mimics of cytoskeletons have been reconstituted in droplets [25, 26] and GUVs [27], capable of e.g. reversible assembly [27, 28] and transport [26]. We recently achieved the formation of contractile DNA rings [29], which may allow morphological dynamics reminiscent of cell division.

Despite these remarkable examples of DNA engineering, there are conceptual challenges related to the use of DNA nanotechnology for the construction of synthetic cells: (i) DNA nanostructures cannot be produced by the synthetic cells themselves. They are first formed by thermal annealing of synthetic oligonucleotides and then encapsulated in lipid vesicles, which is not consistent with metabolism. (ii) Functionality typically requires chemical functionalization of the DNA. DNA nanopores, for instance, were anchored with hydrophobic tags like cholesterol [30]. This further complicates the encoding and production of these structures inside of a vesicle. (iii) Vesicles containing DNA nanostructures have so-far been demonstrated in equilibrium, which is not consistent with *sustaining* life. While chemical fuelling of DNA nanostructures has been demonstrated [31–33], it requires chemical functionalization or DNA strand displacement reactions [34]. (iv) Finally, the implementation of evolution is not straightforward. While it may be possible to genetically encode DNA origami in vesicles [35], purification and annealing steps were still required.

We propose here that these challenges could be addressed using co-transcriptional RNA origami for the construction of synthetic cellular hardware. RNA origami can be genetically encoded in DNA templates [36–38] which can be copied, mutated, evolved, and transcribed using a fraction of the existing machinery from the Central Dogma— only a single protein, namely RNA polymerase. Further, the RNA origami folding mechanism described by Geary *et. al.* in [36, 39] resembles that of proteins—a single polymer chain transcribed by a polymerase undergoes a temporally-segregated series of co-transcriptional folding steps, folding first into local secondary structures (stemloops), followed by the formation of long-distance tertiary structures through internal kissing loops, and finally oligomerization via various quaternary interactions (external kissing loops, overhangs and aptamers). Structures using this folding pathway can be rationally designed using relatively simple heuristics, allowing the creation of a variety of RNA origami structures.

As an initial step toward realizing this RNA-based synthetic cell paradigm, we encapsulated the T7 transcription machinery inside GUVs to facilitate the production of RNA origami from a DNA template (Fig. 1b). To enable continuous feeding of rNTPs and to mitigate waste accumulation often associated with IVTT systems [40], we introduced a pore-forming protein, *α*-hemolysin, on the GUV membrane for the feeding of rNTPs and removal of waste products [41]. Additionally, we implemented another strategy for triggering transcription utilizing a small-molecule ionophore to transport selective ions inside GUVs, facilitating diffusion along concentration gradients while maintaining separation of intracellular and extracellular content [42]. In our pursuit of synthesizing RNA-based cellular structures within synthetic cells, we designed the first 3D RNA origami nanotube compatible with co-transcriptional folding and polymerization in GUVs as mimics of cytoskeletons. While RNA nanotubes have been demonstrated [43, 44], none of them are able to reach micron lengths without thermal annealing. We further show that the small variations in the DNA template sequence can have significant structural and functional impact on the resulting RNA origami nanotubes, establishing genotype-phenotype correlations (Fig. 1c). Finally, we achieved the co-transcriptional functionalization of our RNA-based cytoskeleton with a membrane-binding module, leading to cortex formation and GUV deformation. We envision that the use of genetically encodable functional modules built from RNA may provide a shortcut towards synthetic cells, and the ability to drastically modify the structure and function of RNA-based hardware with small sequence changes will both open new avenues for molecular engineering and shed light on the potential of RNA to perform various cellular functions in an RNA world.

**Fig. 1.**
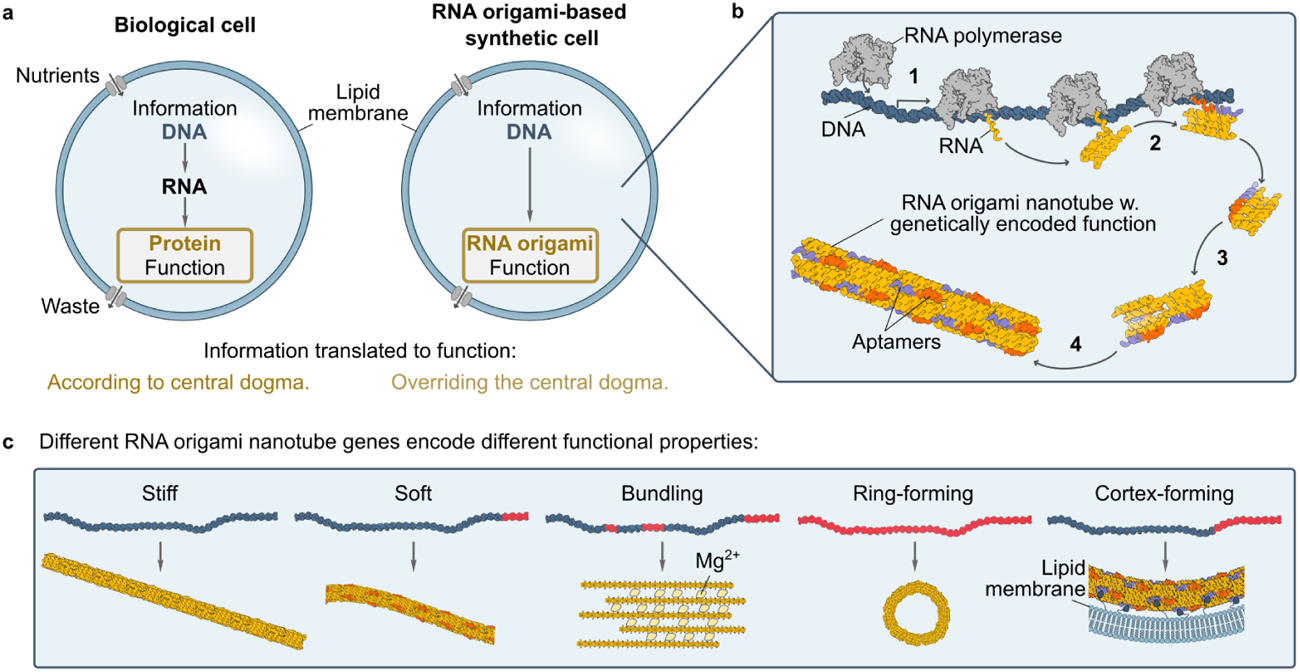
Motivation and conceptualization for engineering RNA origami-based hardware for synthetic cells. **a)** Biological cells function according to the central dogma (DNA serves as a template for RNA, and RNA directs protein synthesis), necessitating the involvement of numerous genes in the translational process. In contrast, a synthetic cell built upon RNA origami requires fewer components while maintaining evolvability. **b)** Mechanism of cotranscriptional RNA origami folding. A DNA template is transcribed by RNA polymerase, whereby the RNA folds up into tiles that self-assemble into higher-order RNA origami nanotubes inside synthetic cells. **c)** Information-function correlation. Mutations on the DNA template result in RNA origami nanotubes with different morphologies and functions.

## 2 Results

### 2.1 Expression of RNA origami inside GUVs

Towards introducing RNA origami as an alternative set of molecular hardware for bottom-up synthetic cell assembly, we first aimed to encapsulate the RNA origami-encoding DNA template and the transcription machinery required for the production of RNA origami inside GUVs as cell-mimicking compartments. We initially chose a previously realized 2D RNA origami which has a well-defined structure and contains a fluorescent iSpinach aptamer attached to the 3’ end (3H-4DT-iSpi, originally designed by [38], see 4.2, Fig. S1a). The RNA origami folds into 3-helix tile subunits, which further assemble into a 5-tile pentagon with 20 nm edges and a width of 7 nm. The folding of the iSpinach aptamer can be detected by measuring the fluoresence of the transcription solution after addition of the fluorophore DFHBI-1T, which increases its brightness more than 800 fold upon binding to the correctly folded aptamer [45, 46]. This allows us to track RNA origami production by measuring fluorescence over time. Compared to *in vitro* transcription (IVT) in bulk, several challenges have to be overcome to realize IVT of RNA origami in GUVs. First, we have to ensure that transcription is indeed happening inside the compartment and not prior to encapsulation. Thus, an external trigger is needed, using a signalling molecule capable of passing the compartment barrier. Second, due to the limited reaction volume of the GUV, nutrient depletion and waste accumulation would stall and favor abortive initiation of the transcription process [47]. To trigger the expression of RNA origami on demand, we introduce two distinct control schemes: restriction of the essential cofactor, Mg^2+^ and restriction of the building blocks, rNTPs.

The semipermeable phospholipid bilayer only allows small hydrophobic and nonpolar molecules to diffuse passively across the membrane [48] while ions like Mg^2+^ and larger charged molecules like NTPs are membrane-impermeable. To achieve highly specific Mg^2+^ import, we chose a Mg^2+^ ionophore [49], which is capable of shuttling Mg^2+^ across the membrane (Fig. 2a).For nucleotide feeding and waste removal, we incorporated the bacterially-derived transmembrane pore *α*-hemolysin in the GUV membrane. With a constriction of 1.4 nm, this pore allows us to continuously fuel transcription with the otherwise membrane-impermeable rNTPs and ions from a bulk feeding solution while preventing the escape of the larger DNA template and product RNA origami structures (Fig. 2b).

To first confirm that Mg^2+^ can indeed be used as a trigger for RNA origami production, we first studied its effect on transcription and folding in bulk. AFM experiments showed that 1 mM Mg^2+^ was insufficient for RNA origami transcription (Fig. 2c,left). On the other hand, if Mg^2+^ concentration was increased from 1 to 6 mM, the transcription and assembly of the pentagonal 3H-4DT-iSpinach structures occurred at high yield (Fig. 2c, right, Fig. 2d, Fig. S1b).

We also tracked successful transcription and folding via a fluorescence assay measuring the successful folding of the fluorescent iSpinach aptamer, which is positioned at the 3’ end of the sequence and thus the last part that is being transcribed (Fig. 2e). In all cases, no fluorescence was observed either in the absence of 6 mM Mg^2+^ or 1 mM rNTPs (empty circles, Fig. 2e), indicating that either Mg^2+^ or rNTPs can be used as a external trigger to control co-transcriptional production of RNA origami. We verified these findings with another RNA origami design, a two-helix tile, named S2T[38], also containing an iSpinach aptamer (Fig. S2).

Next, we demonstrated RNA origami expression in GUVs for the first time. To trigger RNA origami production with Mg^2+^, we loaded the GUVs with the DNA template, rNTPs, all ions along with 1 mM of Mg^2+^ and T7 polymerase during the formation process and added an ionophore and 5 mM Mg^2+^ to the external buffer after GUV formation. We tracked RNA origami production by quantifying iSpinach fluorescence inside of individual GUVs with confocal microscopy for up to three hours (Fig. 2g, Supplementary Movie 2, Fig. S3). The mean fluorescence intensity inside the GUVs increased consistently over time compared to the bulk solution (Fig. 2g), confirming successful RNA origami production inside of the GUVs. We also performed a control where transcription was carried out in the bulk solution, leading to a rapid increase in fluorescence intensity upon the addition of Mg^2+^ to 6 mM (Fig. S4).

For our second control scheme, we reconstituted *α*-hemolysin in the GUVs. This strategy improved the GUV formation efficiency, as all salts required for transcription could also be supplied after formation, which is beneficial because GUV formation is known to be sensitive to high concentrations of divalent ions [50].Next, RNA origami production was effectively initiated by adding rNTPs and tracked inside of individual GUVs (Fig. 2f,h, Supplementary Movie 1), while a control experiment showed similar 3H-4DT-iSpinach transcription patterns triggered with rNTPs in bulk solution (Fig. S4). The results were reproduced faithfully with a second RNA origami design (S2T RNA origami[39], Fig. S5, Supplementary Movie 3). When performing endpoint fluorescence measurements for a larger number of GUVs 4 h or 3 h after adding rNTPs, we observed several fold fluorescence increase for both 3H-4DT-iSpanich and S2T RNA origami (Fig. S6).

**Fig. 2.**
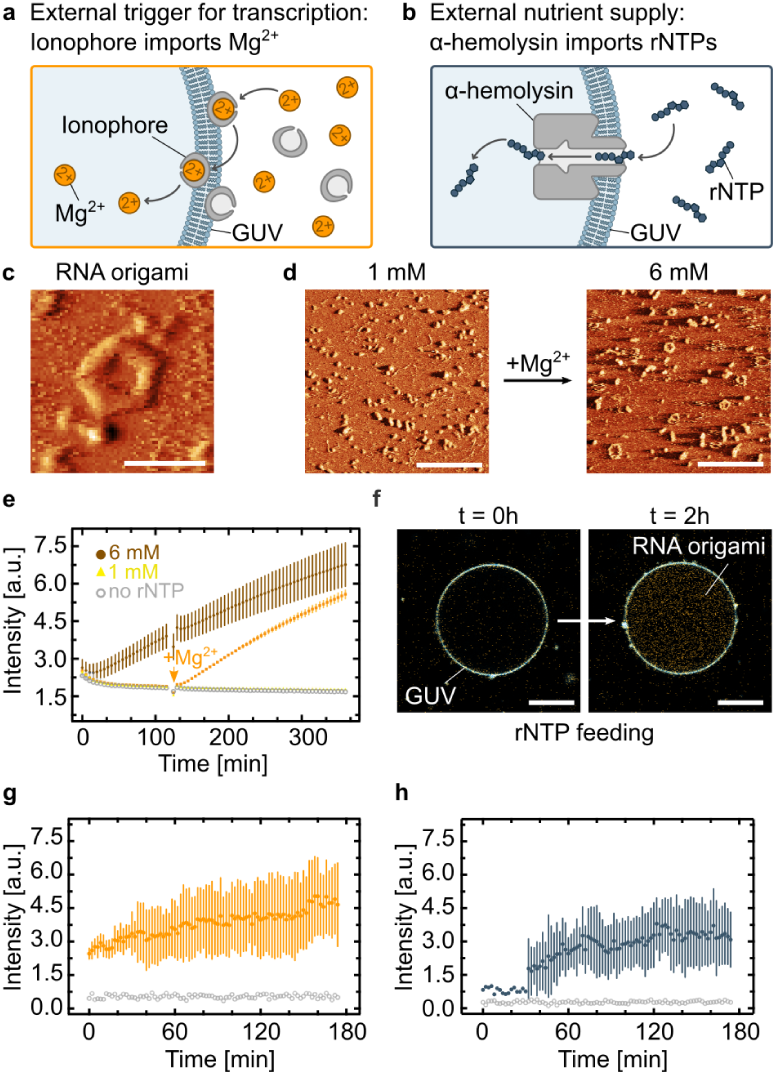
Expression of RNA origami inside GUVs. **a-b)** Schematic illustration showing the external triggers for RNA origami expression by Mg^2+^ import using ionophores **(a)** or rNTP feeding via *α*-hemolysin pores **(b)**. **c)** AFM micrograph of the pentagon-shaped RNA origami (3H-4DT-iSpi), composed of 5 subunits, consisting of three helices each. Scale bar: 50 nm. **d)** AFM micrographs of Mg^2+^-triggered of 3H-4DT-iSpi RNA origami. At 1 mM Mg(OAc)_2_, RNA was incubated for 2 h at mM Mg(OAc)_2_ (left) before transcription trigger with Mg(OAc)_2_ up to 6 mM (incubation time = h, right). Scale bars: 200 nm. **e)** Fluorescence plate reader experiments tracking 3H-4DT-iSpi RNA origami production by measuring DFHBI-1T emission over time (*λex* = 488 nm, mean *±* s.d, *n* = 3 wells). The orange arrow indicates the time point where the Mg(OAc)_2_ concentration was increased from 1 mM to 6 mM in the respective sample. **f)** Confocal overlay images of 3H-4DT-iSpi RNA origami (orange, iSpinach binds DFHBI-1T, *λex* = 488 nm) transcribed inside of a GUV (blue, membrane labelled with DiD, *λex* = 640 nm). Scale bars: 20 µm. **g-h)** RNA origami transcription inside of GUVs triggered by addition of Mg^2+^ (**f**) or rNTPs (**g**) plotted over time (mean *±* s.d, *n* = 6 GUVs). The data was extracted from confocal fluorescence timelapse recordings (Supplementary Movie 1, 2). Empty gray circles denote the background fluorescent signal outside of the GUVs over time.

### 2.2 Construction of RNA origami nanotubes

Having successfully expressed RNA origami in GUVs, we set out to provide a proof of principle for the genetic encoding of RNA origami structures with cell-like functions. Inspired by the success of DNA-based mimics of cytoskeletons [51], we wanted to realize genetically encodable RNA origami nanotubes. For our purposes, an ideal RNA cytoskeleton has to fold cotranscriptionally and should exhibit both cell-sized structures and minimal genetic complexity, with a low number of genes (strands) per functional unit. In line with these criteria, we engineered the first 3D RNA nanotube from a single gene fragment capable of reaching micrometer-scale lengths.

To minimize the number of components, we employed 180° kissing loops for homotypic tile-tile interactions. These external kissing loops allow for more tile-tile cohesion points per strand, which are necessary for strengthening the nanotubes, than traditional sticky-end hybridization methods. Additionally, this approach allows us to compose each tile from a single strand rather than the conventional five strands per tile used in DX tile motifs [44, 52, 53]. Furthermore, unlike traditional multi-strand RNA/DNA nanotube designs where intrinsic curvature is introduced at the tile-tile interaction level using offsets in the helix axes [53], we introduced intrinsic curvature directly at the tile level with dovetail junctions (−3 dovetail) [38, 39]. This produced the desired radial curvature of 120° for a three-tile nanotube where each tile is connected corner-to-corner by 180° kissing loops (Fig. 3a, Fig. S7a). Given that the 11 base-per-turn characteristic of the RNA helix does not enable symmetrical tile designs, we optimized the loop lengths through visual inspection, and generated the sequence using Revolvr [36, 38]. This design process yielded a “wild-type” (WT) tile with approximate dimensions of 11 nm x 5 nm x 2.5 nm and a resultant assembled nanotube diameter of 11 nm (Fig. S7b).

The *in silico* design was subsequently validated *in vitro* by incorporating the sequence into a single DNA template and transcribing it using the same T7 transcription system as before. Successful co-transcriptional folding of RNA origami nanotubes was confirmed with atomic force microscopy (AFM). The nanotubes assembled and reached up to several micrometer in length. AFM yielded an apparent height of the nanotube of 5 to 6 nm which corresponds to at least two RNA duplexes lying on top of one another (an unravelled nanotube would have a height below 2 nm corresponding to a single RNA duplex) (Fig. 3b). At the same time, the measured width is ∼22 nm (Fig. S8a). The deviations from the expected dimensions are thus consistent with the expected tip compression and surface collapse for a hollow nanotube design.

To visualize the RNA origami nanotubes with fluorescence microscopy, we designed a new tile with an iSpinach aptamer extending from the the 3’ end of the tile (hereafter “iSpi”). Again, RNA origami nanotubes were produced co-transcriptionally in one pot under isothermal conditions at 37 *^◦^*C. We obtained cytoskeleton-like networks tens of microns across that could be vizualized with confocal microscopy, likely in association with T7 polymerase due to the co-transcriptional nature of the folding and assembly process (Fig. 3c). Note that compared to the AFM imaging, confocal microscopy was performed without purification. Networks were also observed in AFM if RNA origami nanotubes were not purified from T7 polymerase (Fig. S8b). These are the first reported RNA origami nanotubes that are made of single-stranded tiles in a co-transcriptional manner.

To understand what effect the addition of iSpinach has on the polymerization capacity of the individual tiles, we analyzed the length distribution of the two different variants of the RNA origami nanotubes from AFM images. The *in vitro* transcribed WT nanotubes exhibited the longer continuous segments, with a mean length of 477.4 nm and a 75th percentile length of 585.4 nm. Meanwhile, the iSpi design, whether transcribed in the presence or absence of dye demonstrated mean length below 350 nm, with their 75th percentile lengths falling below 400 nm.

Since the modification of individual tiles is key for the creation of functional cytoskeletons, we need to gain a deeper understanding why modification of the tiles leads to shorter nanotubes. We thus generated a third design modified with a double-stranded loop out overhang (hereafter “dsOV”). This time, we connected the loop out sequence to the tile via a three-way junction (Fig. 3c, for detailed strand layout see Fig. S19). Therefore, the dsOV loop is expected to be connected to the third helix in a less flexible manner (three-way junction) compared to the iSpinach aptamer (three-uracil linker, extended from 3’ end), thus reducing entropic fluctuations which could hinder assembly. Nevertheless, also the dsOV variant of the RNA origami nanotubes were again significantly shorter than WT (Fig. 3d).

To investigate this at molecular resolution, we turned to coarse-grained molecular dynamics (MD) simulations. We simulated each of the three designs as single tiles as well as 1 µm (300-tile) nanotubes using the oxRNA model [54, 55] (Fig. 3e). The 300-tile assemblies were built from relaxed monomers, featuring the largest RNA origami structures ever simulated using oxRNA, containing over 90000 nucleotides (Fig. 3f, Fig. S9). To validate the simulations, we compared the persistence length of the nanotubes extracted from AFM images with the persistence length of the simulated nanotubes (Fig. 3g,h). Experiments yielded a persistence length of 2.7 ± 0.2 µm for the WT and 2.0 ± 0.2 µm for the iSpi design, respectively. In contrast, the dsOV design exhibited a lower persistence length of 1.4 µm. Notably, even though the presence of iSpinach dye, DFHBI-1T, during transcription did not affect the contour length of the RNA origami nanotubes, it significantly reduced the persistence length of the iSpi design by 2.5-fold to 0.8 µm. The MD simulations slightly overestimate the persistence lengths of the RNA origami nanotubes, however, they clearly reproduce the correct order of magnitude and the fact that the dsOV tiles yield the least persistent RNA origami nanotubes. The dsOV tiles show again a drop in persistence length compared to the WT and iSpi designs (1.8-fold and 2.1-fold, respectively).

The ratio of calculated persistence length to mean contour length, which indicates the flexibility of the nanotubes, was approximately 5.6 for both the WT and the iSpi (in absence of dye). This ratio dropped to 2.4 for iSpi in presence of dye and to 4.3 for dsOV design. These results suggest that while the addition of a helix via a single-stranded linker affects the apparent length of the resulting nanotubes (Fig. 3f), it does not impact their flexibility. In contrast, the connection of the overhang helix via a 3-way junction in the dsOV design increases flexibility by 1.3 fold, and the binding of the dye to the aptamer motif increases it by more than 2-fold.

Having confirmed that the simulations reproduce the experimental data sufficiently well, we further examined the molecular-level differences among the design architectures to identify the mechanisms underlying the striking variations in the physical properties of the nanotubes. In the simulations of the dsOV nanotubes, we observe higher rates of bond-breaking throughout the simulation, which are not seen in the WT and iSpi designs (Fig. S10). This indicates that the lower persistence length of dsOV may result from assembly defects and fragmentation of assembled RNA origami nanotubes.

We hypothesize that there are two major factors impacting nanotube stiffness, namely steric hindrance during assembly and intrinsic tile flexibility in the assembled nanotubes. If simply adding an overhang was disrupting nanotube assembly for steric reasons, one would expect the iSpi design to show lower persistence length than the dsOV due to the larger size of the additional structure. However, we do not see this effect, suggesting that the manner of attachment, the simple overhang for the iSpi vs the more rigid 3-way junction for the dsOV, has a significant impact on the ability of the tiles to assemble and the stiffness of the resulting assembly. This hypothesis is further supported by the instability of the dsOV design in oxRNA simulations, where it showed continuous loss of internal bonds over the course of the simulations at a much higher rate than iSpi or WT (Fig. S11). The steric hindrance hypothesis is supported by the difference between the iSpi stiffness with and without the DFHBI dye. When the dye is present and the aptamer is fully structured and fluctuates less, the resulting assemblies are substantially less stiff. The fact that the nanotubes often exhibit kinks rather than smooth contours in AFM ((Fig. S12), also points towards the fact that the primary source of flexibility are broken bonds, transcription errors, or incorrect incorporation of tiles during assembly. This is consistent with the higher persistence lengths measured in simulations, where the nanotubes were first forced to be perfectly formed, as opposed to the stochastic and error-prone polymerization process which preceded the experimental measurements.

It is known that flexibility significantly impacts polymer aggregation. Flexible polymers experience negligible loss of entropy during dimerization, making bundling more favorable compared with semiflexible polymers [56]. Since bundling is key in cytoskeletal organization, we examined our nanotubes for their bundling capacity. We selected the dsOV, which exhibits the highest flexibility without additional ligands/dyes to facilitate the bundling process. As a crosslinker, we chose Mg^2+^, which has been reported as a bundling agent for DNA nanotubes [57].Upon incubation at 20 mM Mg^2+^, comparable cellular Mg^2+^ concentrations, we observed the formation of long nanotube bundles of dsOV design, reaching micrometers in length and a width of 67.91 ± 15.47 nm (Fig. 3i,j, Fig. S13). As expected, bundling was not seen for a semiflexible design like WT (Fig. S14).

**Fig. 3.**
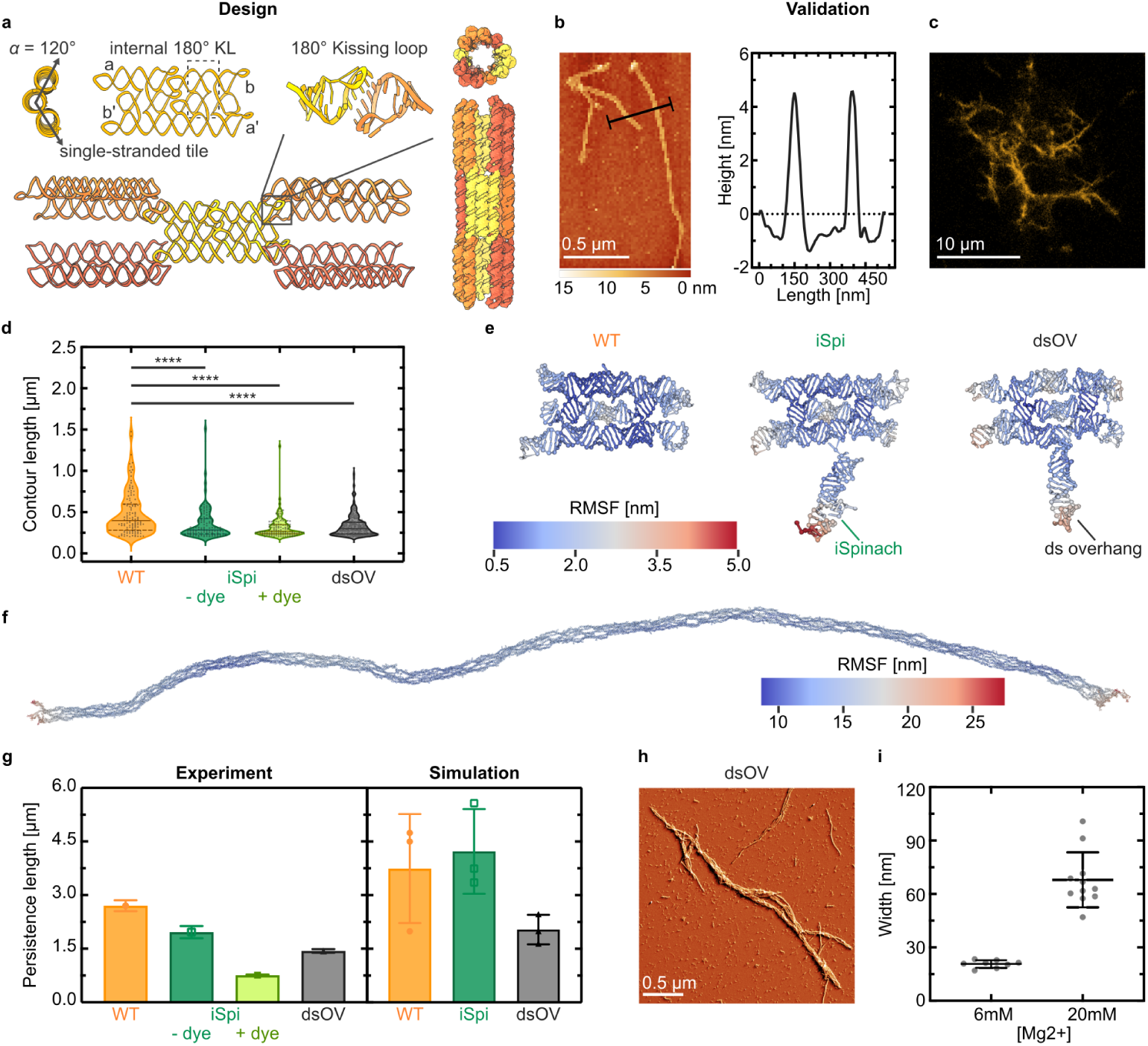
Design and characterization of cytoskeleton-like RNA origami nanotubes. **a)** *In silico* design of RNA origami nanotubes. The nanotube-forming RNA origami tiles assemble co-transcriptionally from a single RNA strand, forming 3 duplexes with *α* = 120° intrinsic curvature. The 180° external kissing loops for nanotube assembly are paired corner-to-corner (a-a’ and b-b’). **b)** AFM micrograph of RNA origami nanotubes produced co-transcriptionally. The height profile is plotted along the black line. **c)** Confocal image of micron-scale, co-transcriptionally folded RNA origami nanotubes after 12 h of IVT. Fluorescence induced by including an iSpinach aptamer and DFHBI-1T dye (*λex* = 482 nm). **d)** Contour length distribution of RNA origami nanotubes extracted from AFM images. The median and quartiles are shown as continuous and dotted lines, respectively. Statistics: two-tailed, parametric, unpaired t-tests with Welch’s correction. Significant (*p <* 0.001) p-values between WT and the other conditions marked with ****. One-way Brown-Forsythe and Welch’s ANOVA tests were performed between the non-WT conditions and both yielded non-significant differences (*p* = 0.0748 and *p* = 0.0494, respectively). Wild-type: WT; iSpinach aptamer on the 3’ end: iSpi, and double-stranded overhang on the 3’ end: dsOV. **e)** OxRNA molecular dynamics simulations of the three monomer designs. **f)** OxRNA molecular dynamics simulation of a 300-tile assembly (here: WT). **e,f)**, show the centroid structure from one simulation run. Each nucleotide is colored by its root-mean-square fluctuation (RMSF). **g)** Persistence length of the RNA origami nanotubes extracted from AFM images (mean *±* s.d error, *n >* 52 nanotubes, see 4.13) and simulations (300-tile nanotubes during the final 10% of the time steps, mean *±* s.d, *n* = 3 simulation runs). **h)** AFM image (error signal mode) of bundled dsOV RNA origami nanotubes upon addition of Mg^2+^ as a crosslinker. The corresponding height image is shown in Fig. S13a. **i)** Width of non-bundled (at 6 mM Mg^2+^) and bundled dsOV RNA origami nanotubes (at 20 mM Mg^2+^) (mean *±* s.d, *n >* 8 nanotubes).

**Fig. 4.**
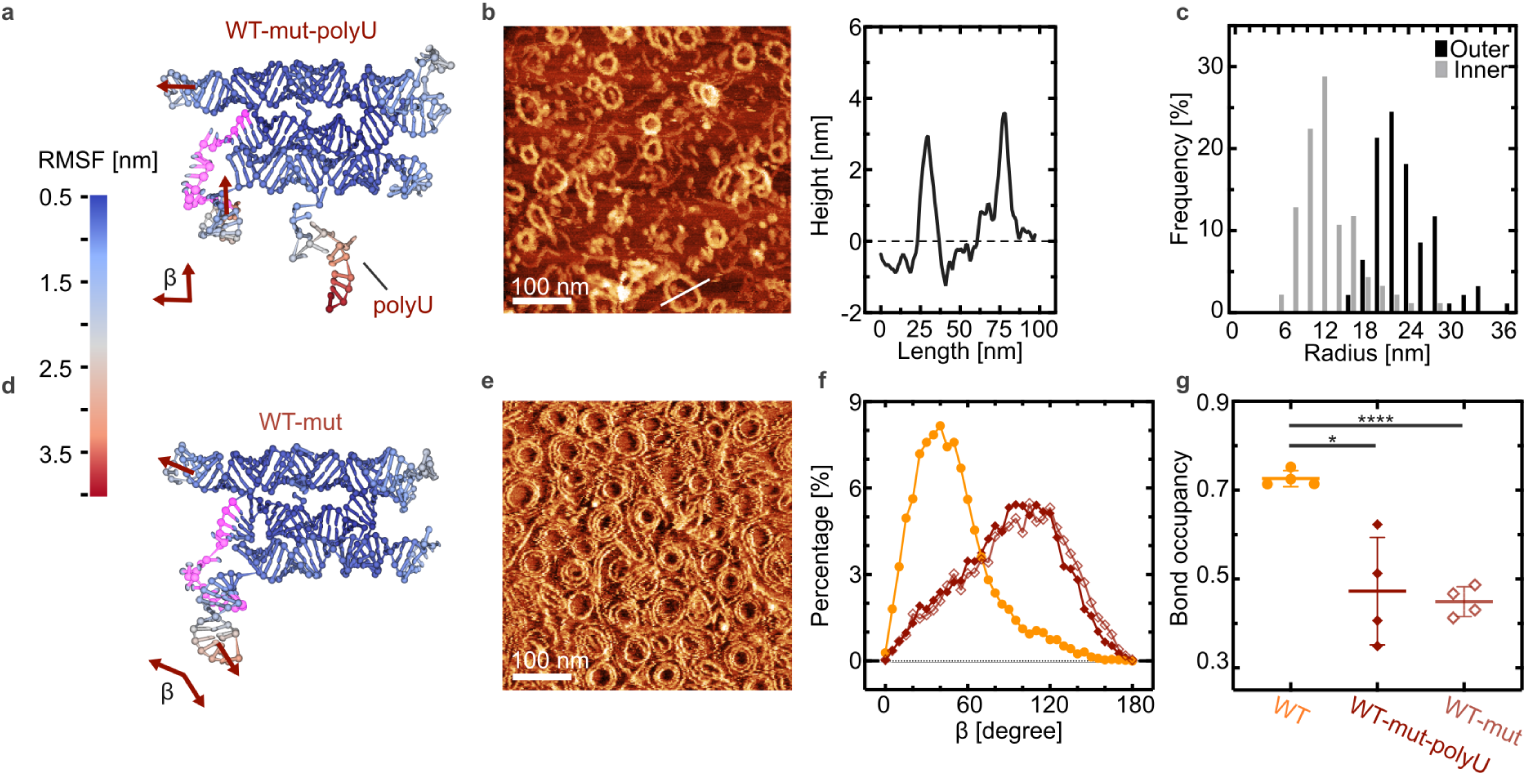
Formation of RNA origami rings. **a,d)** Coarse-grained molecular dynamics simulation of a sequence-mutated WT tile (WT-mut) with (**a**) and without (**d**) single-stranded overhang polyU. *β* denotes the angle between the upper left and lower left duplexes as shown. The broken tetraloop in the middle left helix is highlighted in pink. The centroid structure from one of the simulation runs is shown. Each nucleotide is color coded according to its RMSF. **b)** AFM image and corresponding height profile of *in vitro* transcribed WT-mut-polyU RNA origami rings. The height profile is measured along the white line. **c)** Histogram of the inner and outer radius of RNA origami rings (*n >* 94). **e)** AFM image (lock-in phase channel) of RNA origami rings from the WT-mut tile without overhang. The corresponding height image is shown in Fig. S16b. **f)** Distribution of the angle *β* of simulated single tile using oxRNA coarse-grained simulation (WT: orange circles; WT-mut-polyU: maroon filled diamonds; WT-mut: light-maroon empty diamonds). **g)** Bond occupancy of the tetraloop highlighted in **a,d** (mean *±* s.d, n = 4 simulation runs). A parametric, unpaired t test with Welch’s correction was performed. Two-tailed P values of the comparison between WT and the mutated designs are marked with *** (p *<* 0.001), * (p = 0.0234). The difference is not significant among the mutated designs (p = 0.7295).

### 2.3 Formation of RNA origami rings

During our exploration of RNA origami nanotube designs, we identified a unique structure that consistently formed ring shapes rather than the intended nanotubes. This design used the same strand routing as WT, however with a different sequence produced by Revolvr [38]. It also contained a single-stranded overhang on the 5’ end consisting of the GGAA transcription start sequence and 12 U nucleotides (hereafter termed WT-mut-polyU), intended for binding to a fluorescent probe (Fig. 4a). Unexpectedly, in oxRNA simulations, WT-mut-polyU consistently displayed misfolding in the tetraloop on the “left” side of the middle helix, which was not observed in WT, iSpi or dsOV (Fig. 4a, highlighted sequence, Fig. S15). This region in the new sequence is 50% GC compared to 75% GC composition in the other designs. The nanorings formed both cotranscriptionally (Fig. 4b) and via thermal annealing (Fig. S16a), indicating that the misfolding is inherent to the sequence and not a kinetic trap. Notably, the height of the ring (4 nm, Fig. 4b) was lower than that of the nanotubes (6 nm, Fig. 3b). Characterization of cotranscriptional nanorings revealed an inner and outer radius of 12.77 ± 4 nm and 24.01 ± 4.28 nm, respectively (Fig. 4c), yielding a width of 11.31 nm on AFM.

To confirm that the rings resulted from misfolding in the indicated region, we removed the single-stranded overhang and characterized the resulting assemblies (termed WT-mut). Also without the poly-U, we observed misfolding in the oxRNA simulations of the WT-mut single tiles (Fig. 4d), demonstrating the unbinding is inherent to the sequence of the tetraloop region. In experiments, the WT-mut design without the polyU overhang also resulted in nanorings after both thermal annealing (Fig. 4e) and cotranscriptional, isothermal assembly (Fig. S16b). We characterized the angle *β* between the upper and lower left duplex of the WT-mut tile in the single-tile simulations to assess the effect of misfolding on the connection points among tiles during assembly (Figure.4a,d,e). The WT, WT-mut-polyU and the WT-mut designs exhibited an angle *β* of 49.50 ± 28.82°, 90.61 ± 35.62°, 94.29 ± 37.23°, respectively (Fig. 4f). For nanotube formation, these two duplexes are intended to be parallel to allow for polymerization into a straight nanotube; however, in the mutated designs, the distribution is shifted towards a perpendicular angle between the helices, likely causing a kink that forces the tile to assemble into rings rather than straight tubes. This shift in the *β* angle is likely due to the breakage of the bonds in the misfolded tetraloop region and the corresponding increase in flexibility. Bond occupancy in this region in the single-tile simulations dropped from 0.73 ± 0.02 (WT design) to 0.47 ± 0.12 and 0.45 ± 0.03 for mutated designs with and without the overhang, respectively (Fig. 4g).

### 2.4 Expression of an RNA origami cytoskeleton in synthetic cells

The cytoskeleton plays a crucial role to regulate the cell shape and mechanics. Having built cytoskeleton-like RNA nanotubes that can fold co-transcriptionlly, our next goal was to incorporate these structures into GUVs as synthetic cell models. Production of the iSpi tile was triggered inside GUVs by supplying 1 µM rNPTs externally via reconstituted *α*-hemolysin pores (Supplymentary Movie 4). The production and assembly into cytoskeleton-like structures inside the GUV was monitored with confocal microscopy (Fig. 5a), showing the appearance of RNA origami nanotubes over the course of 6 h. The mean fluorescence intensity inside the GUVs increased severals fold within the first 2 h (Fig. 5b).

We used the area fraction of the RNA fluorescence signal within the GUV to measure the cytoskeleton network growth over time (Fig. 5c, Supplementary Movie 5, 6, 7). Observing cytoskeletal growth within individual GUV is challenging due to the Brownian motion or membrane fluctuation occurring within the 3D compartment (Fig. S17).

Inspired by septin, a natural cytoskeleton component which binds directly to the cell membrane to support membrane organization [58], we aimed for cortex formation as the first function for our RNA cytoskeleton. In order to establish binding between our RNA structures and the vesicle membrane, we added a biotin aptamer to the iSpi tile (Fig. 5d, left, Fig. S19, see 4.2). We then expressed these tiles in GUVs with 5% biotinylated lipids. When transcription was triggered with rNTPs, the RNA nanotubes assembled into a cortex on the inner GUV membrane (Fig. 5d, right, Supplementary Movie 8). RNA origami nanotubes were exclusively observed bound to the biotinylated membrane in presence of biotin aptamer (Fig. S17b), whereas without the biotin aptamer, RNA origami nanotubes were found in the GUV lumen as in previous experiments (Fig. S17b). The membrane binding function results in a shift in the center of mass of the RNA fluorescence signal towards synthetic cell membrane (Fig. 5e, Fig. S17d,e). Interestingly, we occassionally observed that, like septin [58], biotin aptamer-based RNA structures are capable of promoting GUV deformation and inducing membrane curvature (Fig. 5f).

**Fig. 5.**
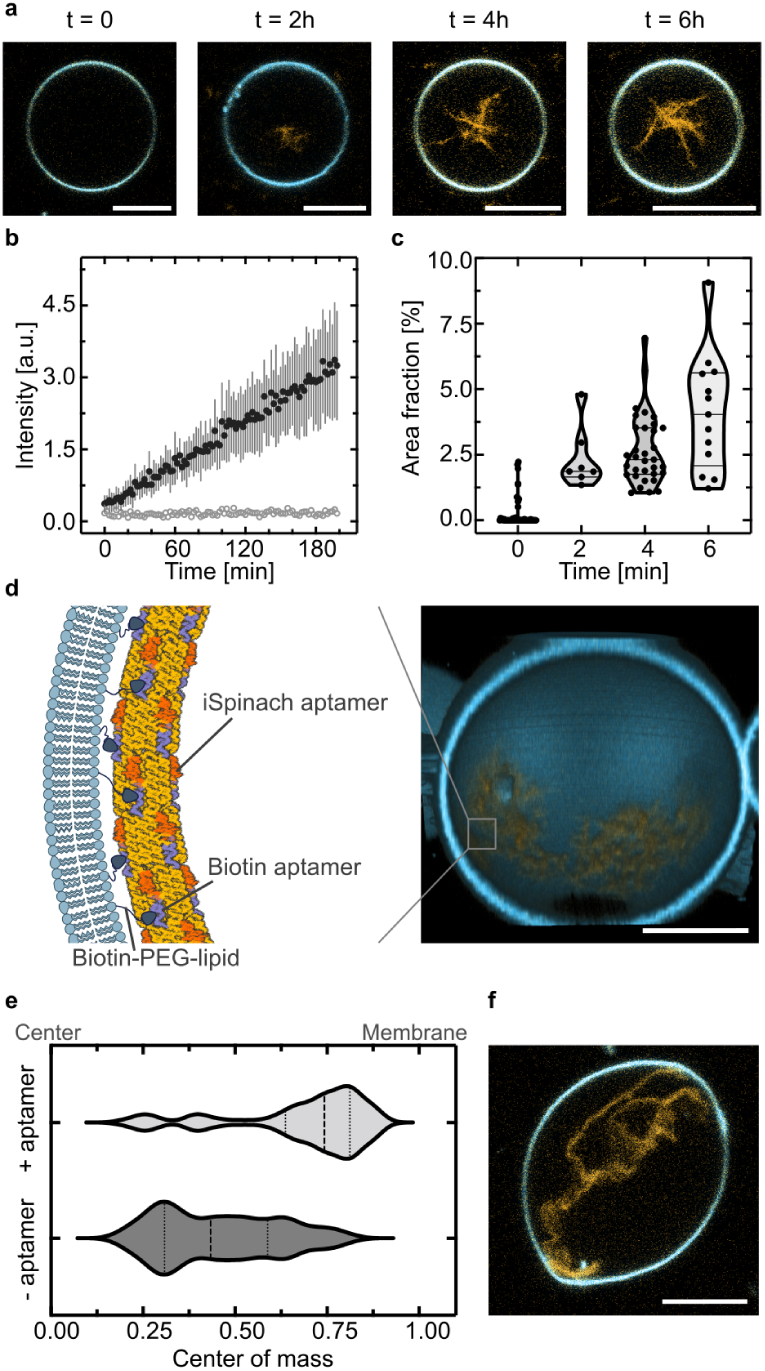
Expression of RNA origami cytoskeletons in synthetic cells. **a)** Confocal time series of cytoskeleton-like iSpi RNA origami nanotubes (orange, *λex* = 482 nm) expressed inside a GUV (blue, membrane labelled with DiD, *λex* = 644 nm). Scale bars: 10 µm. **b)** RNA nanotube transcription inside of GUVs triggered by addition of rNTPs plotted over the first 3 h of expression (mean *±* s.d, *n* = 6 GUVs). The data was extracted from confocal fluorescence timelapse recordings (Supplementary Movie 4). Background fluorescence outside of the GUVs is plotted as gray circles. **c)** Area fraction occupied by the RNA origami nanotubes plotted over time (*n* = 7). **d)** Cortex formation with RNA origami nanotubes on the inner GUV membrane. Left: Schematic representation of an RNA origami nanotube adhering to the biotinylated GUV membrane via a biotin aptamer attached to the iSpi tile. Right: Confocal 3D reconstruction of an RNA origami cortex on the inner GUV membrane after its expression. **e)** Distribution of RNA nanotubes with or without the biotin aptamer inside of the GUV. The distribution of the center of mass of the iSpi fluorescence relative to the GUV center is plotted (*n >* 78 GUVs, median: dashed lines, first and third quartile: dotted lines. **f)** GUV deformation caused by biotin aptamer-functionalized RNA origami nanotubes. Scale bars: 10 µm.

## 3 Conclusion

In conclusion, we have provided a conceptual framework for the construction of synthetic cells based on the expression of RNA origami inside of lipid vesicles. It offers advantages compared to both DNA origami as well as proteins as molecular hardware in synthetic cells. Compared to DNA origami, RNA is genetically encodable, which means that the synthetic cells can produce their own building blocks in a metabolic out-of-equilibrium process. This makes the approach different from the mere encapsulation of DNA origami structures in vesicles. Additionally, we have shown that we can incorporate functionality like membrane binding by using RNA aptamers instead of covalent modifications needed on DNA structures. Finally, subtle changes to the design, like simply swapping out overhanging sequences or weakening specific helices, can result in substantial morphological changes, hinting towards exciting prospects for directed evolution approaches. Compared to proteins, expression in GUVs required far fewer components— only T7 polymerase compared to the entire transcription-translation machinery. This could mean that the realization of a self-sustained chemical system capable of evolution according to NASA’s definition of life becomes more accessible [59, 60]—although this remains speculative for now and diverse scientific communities may, one day, realize diverse variants of synthetic cells. From a practical point of view and beyond synthetic life, genetically encodable hardware made from RNA origami may lead to new exciting biophysical tools to manipulate cells and to address questions in cell biology.

Our first use of RNA origami in the context of synthetic cells is just a start. In the future, we expect RNA origami-based hardware to incorporate ribozymes, i.e. catalytically active RNA sequences, for the realization of molecular machines which execute complex cellular tasks. From a more fundamental point of view, the study of ribozyme activity in confinement has been shown to deviate from bulk activity [61] [62], which hints towards exciting prospects. Additionally, it will be interesting to look for ways to reproduce the DNA template, e.g. by templated ligation [63], and the use of polymerase ribozymes [64] instead of T7 for RNA production or for the direct self-replication of the RNA origami. Notably, the construction of synthetic cells based on co-transcriptional RNA origami is inherently compatible with evolution and AI-guided design, which allows orders of magnitude higher throughput and design testing compared to traditional rational engineering techniques, providing solutions well beyond known structure-function relationships. This will increase the functional complexity of synthetic cells, while at the same time, incorporating evolution, as a fundamental characteristic of life, into the design process itself. We may be curious to find out if an RNA version of synthetic life may also provide perspectives for the origins of life in the context of the “RNA world”.

## 4 Methods

### 4.1 Materials

DOPC(1,2-dioleoyl-*sn*-glycero-3-phosphocholine, cat *#*850375), DOPG(1,2-dioleoyl-*sn*-glycero-3-phospho-(1’-*rac*-glycerol) (sodium salt), (cat *#*840475) in chloroform were purchased from Avanti Polar Lipids. The membrane dye DiD (1,1’-Dioctadecyl-3,3,3’,3’-Tetramethylindodicarbocyanine, 4-Chlorobenzenesulfonate salt, cat *#*D7757) and T7 RNA polymerase (cat *#*EP0112) were purchased from Thermo-Fisher Scientific. Sodium acetate (cat *#*S2889), Potassium chloride (cat *#*P3911), Sodium hydroxide (cat *#*S5881), Polyvinyl alcohol (cat *#*S8062894), DL-Dithiothreitol (cat *#*DO632), Sucrose (cat *#*S0389), Tris-acetate (cat *#*T1258), Glucose (cat *#*G6152), Magnesium-Ionophor I (N,N’-Diheptyl-N,N’-dimethyl-1,4-butandiamid)(cat *#*63082) and *α*-hemolysin from Staphylococcus aureus (cat *#*H9395) were purchased from Sigma-Aldrich. Magnesium acetate tetrahydrate (cat *#*A322119) was purchased from Merck. Phusion® High-Fidelity PCR Kit (cat *#*E0553S) and RNase Inhibitor, Murine (cat *#*M0314) were purchased from New England Biolabs. Nuclease free water was purchased from Integrated DNA Technologies. Microscopy experiments were performed in 18-well 15 *µ*-slide glass bottom chambers (cat *#*81817) from ibidi.

### 4.2 Design of RNA origami

The design of the RNA tiles was guided by the principles established by Geary et al. for ssRNA origami [39].

The 3H-4DT-iSpinach blueprint was generated by adding the iSpinach module to the 3’ end of the 3H-4DT design from Geary et al. [38]. The iSpinach aptamer was connected to the core design by a 2 Uracil linker.

The blueprints for the nanotube-forming tiles were designed based on the 3H-3DT tile from Geary et al. [38]. The first and third helices were extended so that the helical turn of the external 180°kissing loops would align upon assembly. This extension process was done manually via cycles of extension and visualization using the RNAbuild script from ROAD[38] and ChimeraX [65] to minimize the distance between kissing loops upon closure of the tube. During this process, the length of the helices were also adjusted so that the internal kissing loops would correspond to 8 base pair duplexes instead of the original 9 in the 3H-3DT design, reflecting the updated understanding of internal kissing loop structure based on cryoEM structures [66].

The blueprint of iSpi design was then put into Revolvr[38]. The final sequence was chosen using two criteria: both external kissing loop pairs should have binding energy below −8 kcal mol*^−^*^1^; and the difference in binding energy between the two pairs should be minimal. The WT design was created by removing the iSpi overhang. The dsOV was introduced into the tile blueprint without further sequence optimization. All blueprints were used as input for the trace analysis script from ROAD [38] to check for folding irregularities and then used for oxRNA simulations.

The blueprints of the mutated ring-forming tiles were generated as described in 2.3 and the sequence optimization was performed in the same manner described for the other tiles.

The RNA origami containing biotin aptamer was designed by concatenating the WT, biotin aptamer and iSpinach aptamer in the following sequence: WT tile-AAA-biotin aptamer-AAAA-iSpinach aptamer (Fig. S19). The sequence was then checked using RNAfold Web Server to ensure correct folding of each aptamer in the concatenated sequence.

### 4.3 Synthesis of RNA origami in bulk

DNA templates were synthesized as double-stranded gBlocks from Integrated DNA Technologies (IDT). DNA gBlocks (0.5 ng µL*^−^*^1^) were PCR-amplified using 14–25-nt primers (See Supplementary Data 1), using Phusion^©^ High-Fidelity PCR Kit (NEB) with annealing temperature at 62 *^◦^*C. The PCR product was then purified using a Qiagen PCR purification kit. RNA was transcribed and cotranscriptionally folded in a one-pot reaction containing PCR purified DNA template (4 ng µL*^−^*^1^), *Mg*(*OAc*)_2_ (6 mM), NaOAc (40 mM), KCl 40 mM, Tris-OAc (50 mM, pH 7.8), rNTPs (1 mM each), dithiothreitol (DTT) (1 mM), and DFHBI-1T dye (62.5 µM) and RNase inhibitor (1 U µL*^−^*^1^), if not described otherwise. Reactions were initiated by adding T7 RNA polymerase (0.2 U/*µ*l). Transcription reactions were carried out in 100-*µ*l volumes at 37 *^◦^*C for 2 h to 12 h, depending on the sequence length. Nanotube designs required longer transcription times (4-12 h). All the experiments were performed using nuclease free water.

### 4.4 Atomic force microscopy

RNA origami solution was pre-diluted to avoid structure overlapping on the mica if necessary. Subsequently, 20 µL of the solution mixed with 20 µL of imaging buffer (40 mM Tris-Acetate containing 1 mM EDTA and 20 mM Mg ^2+^) were deposited onto a freshly cleaved mica surface (0.95 cm diameter, (Science Services GmbH, Munich, Germany) and allowed to adsorb for a minimum of 1 min. The liquid chamber was then filled with 1 mL of imaging buffer. The sample was imaged using Nanowizard 2 high-speed atomic force microscope (Bruker) in either AC fast imaging mode or QI^TM^ mode with a JPK Nanowizard ULTRA Speed using FASTSCAN-D cantilevers (f = 110 kHz, k = 0.25 N m*^−^*^1^, Bruker Nano Inc., Camarillo, CA, USA).

### 4.5 Fluorescence assay in bulk

3H-4DT RNA origami with an iSpinach fluorophore was prepared with PCR purified DNA template (4 ng µL*^−^*^1^), NaOAc (40 mM), KCl (40 mM), Tris-OAc (50 mM, pH 7.8), rNTPs (1 mM each), dithiothreitol (DTT)(1 mM), DFHBI-1T dye (62.5 µM) in four different conditions of Mg(OAc)_2_ (*i*.e. 0 mM, 1 mM, 1 to 6 mM, 6 mM). In 1 to 6 mM sample, first 1 mM Mg(OAc)_2_ was added for 2 h and then up to 5 mMof Mg(OAc)_2_ added for next 2 h. To measure the fluorescence intensity, 100 µLfinal volume was pipetted into a Greiner 96-well black bottom plate (Sigma-Aldrich) which was finally placed inside the plate reader (Spark multimode plate reader from Tecan Life Science) for 4 h at 37 *^◦^*C. RNA production was quanified by measuring the emission of DFHBI-1T upon excitation with 488 nm (DFHBI-1T dye: *λ_ex_*= 482 nm, *λ_em_* = 505 nm) with a gain of 100 (manually set).

### 4.6 GUV preparation

GUVs were prepared by the polyvinylalcohol (PVA)-gel assisted swelling method[41]. In detail, a PVA solution was prepared by mixing 5% (w/v) PVA (MW: 145 000 Da) in nuclease free water with sucrose (100 mM) for 24 h, at 400 rpm, at 90 *^◦^*C. The PVA solution (50 µL) was dried as a thin film on a glass slide (60 mm x 24 mm) at 50 *^◦^*Cfor 30 min. Then, 5 µL of a lipid mixture in chloroform containing 10 mol% DOPG (10 µg µL*^−^*^1^) and 1 mol% DiD dye in 10 µg µL DOPC was spread onto the PVA layer and dried for 1 h at 37 *^◦^*C. The DiD dye stock solution (10 µg µL*^−^*^1^) was prepared in chloroform. Using a Teflon chamber (ca. 40 mm x 24 mm) as a spacer and a second glass slide, a chamber was assembled on top of the slide with the lipid-coated PVA. Then, the lipids were hydrated with 1 mL of 100 mM sucrose containing all the transcription components to be loaded into the GUV depending on the experiment for 1 h at room temperature, allowing for GUV formation. After that, the chamber was inverted for 5 min, gently taped twice using a pipette tip and the GUVs were harvested into a 1.5 mL Eppendorf tube and left to settle for 30 min. For the washing, 350 µL of GUVs was taken from bottom and added 1 mL of 150 mM glucose buffer and the GUVs were allowed to settle overnight at 4 *^◦^*C. The next day morning, the top 1 mL of the buffer was removed without disturbing the bottom layer. The GUVs were washed second time with 0.5 mL of 150 mM glucose buffer and the GUVs were only allowed to settle for 2 to 3 h at 4 *^◦^*C. For subsequent experiments, the GUV solution was sourced from the bottom, where there was a higher likelihood of obtaining GUVs with effectively encapsulated DNA templates and RNA polymerase, attributable to their greater density. All experiments were performed using nuclease free water. For imaging, an 18-well ibidi glass bottom chamber was pre-incubated with 3% BSA solution and washed with nuclease free water twice before GUV addition.

### 4.7 Confocal Microscopy

All images were acquired on a LSM 900 confocal laser scanning microscope from Carl Zeiss AG, Germany equipped with two excitation lasers (488 nm and 640 nm). For the live observation of the RNA origami expression (Figure 2), the 20x air objective was used. Expression of RNA origami nanotubes was imaged using a 63X water objective. The GUV membrane was labelled with DiD (*λ_ex_* = 644 nm, *λ_em_*= 665 nm)(Sigma-Aldrich). The RNA origami contained an iSpinach aptamer which binds the fluorophore DFHBI-1T (*λ_ex_* = 482 nm, *λ_em_*= 505 nm.

### 4.8 Expression of RNA origami in GUVs triggered by Mg^2+^

GUVs containing a mixture of the DNA template (4 ng µL*^−^*^1^), RNA polymerase (0.2 U/*µ*L), each rNTP (1 mM), Mg(OAc)_2_ (1 mM), NaOAc (40 mM), KCl (40 mM), Tris-OAc (pH 7.8, 50 mM), DTT (1 mM), DFHBI-1T dye (62.5 µM) and sucrose (150 mM) were formed as described. Subsequently, a specific washing protocol was employed, wherein 350 µL of GUVs were rinsed once with 1 mL of 570 mM glucose buffer, maintained at room temperature for 2 to 3 h. On the same day, 100 µL of the washed GUV solution was transferred to an 18-well imaging chamber and supplemented with 10 mM Magnesium-Ionophor I (Merck). Once the sample was placed on the confocal microscope with an incubation chamber held at 37 *^◦^*C for 2 h. Following this incubation, an additional 5 mM of Mg(OAc)_2_ was introduced into the GUV solution. The GUVs were continuously monitored for up to four hours.

### 4.9 Expression of RNA origami triggered by rNTPs in GUVs

GUVs containing a mixture of the DNA template (4 ng µL), RNA polymerase (0.2 U/*µ*L), *α*-hemolysin (15 ng µL) and sucrose (100 mM) were formed as described. On the following day, 80 µL of purified GUVs were supplemented with a feeding solution containing Mg(OAc)_2_ (6 mM), NaOAc (40 mM), KCl (40 mM), Tris-OAc (pH 7.8, 50 mM), DTT (1 mM)and DFHBI-1T dye (62.5 µM). These components could enter the GUV lumen via the *α*-hemolysin pores. The GUVs, now containing the necessary components for co-transcriptional folding except rNTPs for 2 h at 37 *^◦^*C, were transferred to an 18-well imaging chamber and allowed to settle to the bottom of the slide for 20 min before imaging commenced. Once the sample was placed on the confocal microscope with an incubation chamber held at 37 *^◦^*C, rNTPs were added externally to achieve a final concentration of 1 mM for each nucleotide. The rNTPs also translocate into the GUVs via *α*-hemolysin. GUVs were monitored over a period of up to four hours. Alternatively, to image the formation of the RNA origami cytoskeleton-like at discrete time points (0 h, 2 h, 4 h, and 6 h), GUVs were incubated with all above mentioned components including rNTPs in an Eppendorf tube within a thermal block set to 37 *^◦^*C prior to imaging.

### 4.10 Analysis of RNA origami expression in GUV

To quantify the fluorescence intensity within GUVs where transcription was triggered with Mg^2+^ or rNTPs (as shown in Fig. 2f, 2g, 5b, S5b, and S6), GUVs were selected using the DiD channel (*λ_ex_* = 644 nm, *λ_em_* = 665 nm). The fluorescence intensity was then analyzed in the DFHBI-1T channel (*λ_ex_* = 482 nm, *λ_em_*= 505 nm). A region of interest (ROI) for each GUV was delineated with a circular mask on an 8-bit grayscale image using Image J (1.53v). To analyse the transcription trigger over time, the same GUV timeline image (as shown in Supplymentary movie 1, 2, 3, 4) was used at 2 min interval up to 3 h to 4 h. The mean intensity values obtained for each GUV were obtained by using ‘Measure’ function and presented as fluorescence intensity across the samples.

### 4.11 Purification of RNA origami

RNA origami was purified using centrifugal filters with a pore size of 100 kDa (Merck Millipore). After centrifugation for 5 min at 10 000 g, 1x PBS (pH 7.4) containing 20mM MgCl_2_ was added so that the final solution is 2× concentrated. The purified solution was then used for AFM imaging as described above.

### 4.12 oxRNA simulations

Coarse-grained modeling of individual RNA tiles and the nanotubes was performed using the oxRNA2 force field [54, 67] with the CUDA-accelerated oxDNA molecular dynamics (MD) simulation engine (version 3.5.2) [55, 68, 69]. Structures were exported from ROAD schematics to PDB format using the RNAbuild script from ROAD [38]. The PDB structures were then converted to oxDNA simulation files using a local copy of TacoxDNA updated to work with Python 3.11 [70] (pipeline script in 5). Structures were relaxed using the protocol detailed in [71, 72]. Briefly, a short Monte-Carlo simulation was performed to remove excluded volume clashes between nucleotides, followed by a longer (at least 5e7 steps, *dt* = 0.003) MD relaxation performed using the “Langevin” thermostat and a modified backbone FENE potential to avoid numerical instabilities due to high forces. Harmonic traps were placed between all paired nucleotides to maintain the structure during relaxation. Successful relaxation was verified by observing per-nucleotide energy values below -1.4 and visualization of the intended structure with oxView [72, 73].

Multiple production equilibrium MD runs were performed on NVIDIA a100 GPUs using edge-based calculation parallelization [69] for 1e9 simulation steps at *dt* = 0.003. This roughly corresponds to a timestep of 9.09 fs and a total simulation time of 9.09 µs by direct unit conversion. It should be noted, however, that calculating exact time correspondence in coarse-grained simulations is not straightforward. Based on a comparison between DNA binding rates in experiments and oxDNA simulations (see the SI of [74]), the time represented by these simulations may be closer to 30 ms. The temperature was maintained at 37 *^◦^*C using an Andersen-like thermostat (thermostat = “brownian” or “john” in the oxDNA parameter file) [75]. Debye-Huckel electrostatics were applied to the system mimicking 0.5 M NaCl [67]. Configurations were saved every 5e5 steps, resulting in trajectories with 2000 frames each for analysis. Since the dynamics of correctly folded structures are of interest, we ran multiple simulations and selected the first four simulations where the internal kissing loops maintained a minimum of 11 out of designed 12 bonds on average throughout the simulation for further analysis.

The mean structures, RMSFs, and bond occupancies were calculated from simulation trajectories using oxDNA Analysis Tools (OAT, version 2.3.5) [73]. The structure closest to the mean structure was extracted from one replicate for each structure using the centroid function in OAT and this structure used to build the nanotube simulations using a custom-written Python script (See 5), we built 100-layer (300 tiles, ∼80,000 nucleotides) simulations from each single tile. The nanotubes were then relaxed with externally-applied harmonic traps to ensure that the kissing loop cohesion was successfully formed. Once over 90% of target bonds were formed, the forces were dropped and triplicate production simulations with the same parameters as the single tiles were performed.

### 4.13 Persistence length analysis

All AFM images were pre-processed with the open-access software Gwydion [76] using standardized Align Rows and Levelling tools, then exported into Tagged Image File Format (TIFF). The TIFF images were then analyzed with ImageJ2 (ImageJ2 2.14.0/1.54f). The images were cropped manually to separate individual nanotubes. The nanotube detection was done automatically on the cropped images using a custom-written ImageJ macro script (See 5). In brief, the script used Ridge Detection plugin [77, 78] and adjusted the parameters according to the image histogram and scale. The minimum line length and line width were set to 107.4 nm and 17.1 nm, respectively.

The coordinates extracted from ImageJ were then used as input for a custom-written Python script (See 5) to calculate the persistence length of the *in vitro* transcribed nanotubes. To reduce detection error due to low resolution in imaging, the data was filtered to exclude all detection under 200 nm. The squared end-to-end distance of each nanotube was plotted against its contour length. The data was then fit using the theoretical relation:

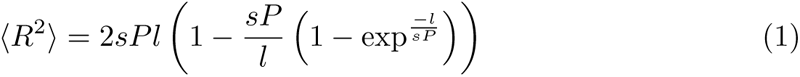

where *R*^2^ is the squared end-to-end distance, *l* is the contour length, *P* is the persistence length and *s* is a surface parameter that is set to 1. For each nanotube, the end-to-end distance and the total contour length of the nanotube is extracted for fitting. The data is masked by residuals ±1 standard deviation. For the simulated structures, due to the narrow distribution of contour length (as to the simulation is only one nanotube over time), one persistence length is calculated for each time point. The end-to-end distance of every pair of points and their contour length are fitted using the same equation and masking as described above. The data plotted in Fig. 3 is the last 10% of the simulation frames (*t* ≥ 9e8).

### 4.14 RNA origami bundling

RNA was transcribed *in vitro* for 2 h. Subsequently the samples were heated to 75 *^◦^*C for 1 min and then cooled down to 25 *^◦^*C with a temperature gradient of −0.5 *^◦^*C per min in a thermocycler (BioRad). For bundling, 1 µL of annealed RNA origami was mixed with 20 µL of imaging buffer (40 mM Tris-Acetate containing 1 mM EDTA and 20 mM Mg ^2+^) and deposited onto a freshly cleaved mica to adsorb for 5 min. The AFM images were processed using Gwyddion. Five width measurements, equally distributed along the measured object, were performed and the averaged widths were reported in Fig. 3j.

### 4.15 Analysis of RNA origami nanotube growth

Images of nanotube-producing GUVs at different time points (Fig. 5c) were analysed in ImageJ2 (ImageJ2 2.14.0/1.54f) using custom-written ImageJ macro scripts (See 5). In summary, for time point t = 0 h, the GUVs were automatically detected using the Hough Circle Transform plugin. For further time points, the GUVs which produced RNA nanotubes were manually chosen. The regions of interest in the RNA fluorescence channel were then filtered and threshold using Gaussian Blur, Convert to Mask and Erode functions. The area fraction was then extracted for each GUV using Measure function.

### 4.16 Analysis of RNA cytoskeleton attachment to GUV membranes

Images of single GUVs that produced RNA cytoskeleton with and without biotin aptamer were analysed in ImageJ2 (ImageJ2 2.14.0/1.54f) using custom-written ImageJ macro scripts (See 5). The GUVs were automatically detected using Analyze Particles method. In short, the lipid fluorescence channel was processed using Auto Threshold, Fill Holes, Erode, Gaussian Blur then Convert to Mask functions prior to particle analysis. The detected regions of interest were then manually filtered to remove false detection of non-GUV particles. The radial profile of each selected region of interest was extracted using the Radial Profile function. The origin of the radial profile was the measured centroid of the region of interest. The radius is the primary axis of a fitted ellipse to the region of interest.

The radial profile was then analyzed and plotted using a custom-written Python script (See 5). The center of mass of the RNA fluorescence relative to the GUV center was calculated for each GUV using the formula:

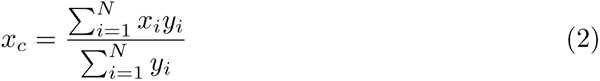

where *x_c_* is the center of mass of the radial fluorescence, *x_i_* are the normalized distance from the center of the GUV, *y_i_* is the normalized RNA radial fluorescence at distance *x_i_*, and *N* is the number of *x*, *y* pairs.

### 4.17 Data visualization and analysis

Plot graphing and statistical tests were performed using GraphPad Prism (Version 5 and Version 10.2.3).

## 5 Code availability

All custom written code was deposited on GitHub at [available upon acceptance]. https://github.com/maiptran21/RNAcytoskeleton

## 6 Data availability

Supporting data is available in the Supplementary Information. All raw data and analysis files used in the study are deposited on HeiData (doi: available upon acceptance).

## Acknowledgements

K.G. acknowledges funding from the ERC starting grant ENSYNC (101076997). M.P.T. and K.G. thank the Hector Fellow Academy. M.P.T. and K.G. received funding by the Federal Ministry of Education and Research (BMBF) and the Ministry of Science Baden-Württemberg within the framework of the Excellence Strategy of the Federal and State Governments of Germany. E.P. and T.C. were supported through state funds approved by the State Parliament of Baden-Württemberg for the Innovation Campus Health + Life Science Alliance Heidelberg Mannheim. MD simulations were performed on the HPC system Raven at the Max Planck Computing and Data Facility. All authors thank Claudia Helbig for helpful technical support and troubleshooting.

## References

[1] Oskar Staufer, O., et al. Building a community to engineer synthetic cells and organelles from the bottom-up. eLife 10, e73556 (2021).

[2] Crick, F. H. C. On protein synthesis, Vol. 12, 138–163 (The Society for Experimental Biology, 1958).

[3] De Capitani, J. & Mutschler, H. The long road to a synthetic self-replicating central dogma. Biochemistry 62, 1221–1232 (2023).

[4] Vogele, K. et al. Towards synthetic cells using peptide-based reaction compartments. Nature Communications 9, 3862 (2018).

[5] Blanken, D., Foschepoth, D., Serrão, A. C. & Danelon, C. Genetically controlled membrane synthesis in liposomes. Nature communications 11, 4317 (2020).

[6] Xu, C., Martin, N., Li, M. & Mann, S. Living material assembly of bacteriogenic protocells. Nature 609, 1029–1037 (2022).

[7] Berhanu, S., Ueda, T. & Kuruma, Y. Artificial photosynthetic cell producing energy for protein synthesis. Nature communications 10, 1325 (2019).

[8] Garenne, D. et al. Cell-free gene expression. Nature Reviews Methods Primers 1, 49 (2021).

[9] Forster, A. C. & Church, G. M. Towards synthesis of a minimal cell. Molecular systems biology 2, 45 (2006).

[10] Lavickova, B., Laohakunakorn, N. & Maerkl, S. J. A partially self-regenerating synthetic cell. Nature Communications 11, 6340 (2020).

[11] Robertson, M. P. & Joyce, G. F. The origins of the rna world. Cold Spring Harbor perspectives in biology 4, a003608 (2012).

[12] Rothemund, P. W. Folding dna to create nanoscale shapes and patterns. Nature 440, 297–302 (2006).

[13] Göpfrich, K., Platzman, I. & Spatz, J. P. Mastering Complexity: Towards Bottom-up Construction of Multifunctional Eukaryotic Synthetic Cells. Trends in Biotechnology 36, 938–951 (2018).

[14] Luo, L. et al. Dna nanopores as artificial membrane channels for bioprotonics. Nature Communications 14, 5364 (2023).

[15] Göpfrich, K. & Keyser, U. F. Dna nanotechnology for building sensors, nanopores and ion-channels. Biological and Bio-inspired Nanomaterials: Properties and Assembly Mechanisms 331–370 (2019).

[16] Jahnke, K. et al. Dna origami signaling units transduce chemical and mechanical signals in synthetic cells. Advanced Functional Materials 34, 2301176 (2024).

[17] Marsden, H. R., Korobko, A. V., Zheng, T., Voskuhl, J. & Kros, A. Controlled liposome fusion mediated by snare protein mimics. Biomaterials science 1, 1046– 1054 (2013).

[18] Adamala, K. P., Martin-Alarcon, D. A., Guthrie-Honea, K. R. & Boyden, E. S. Engineering genetic circuit interactions within and between synthetic minimal cells. Nature chemistry 9, 431–439 (2017).

[19] Simunovic, M. et al. How curvature-generating proteins build scaffolds on membrane nanotubes. Proceedings of the National Academy of Sciences 113, 11226–11231 (2016).

[20] Xiang, Y., Lyu, R. & Hu, J. Oligomeric scaffolding for curvature generation by er tubule-forming proteins. nature communications 14, 2617 (2023).

[21] Hadorn, M. & Eggenberger Hotz, P. Dna-mediated self-assembly of artificial vesicles. PLoS One 5, e9886 (2010).

[22] Fu, Y.-H. et al. Constructing artificial gap junctions to mediate intercellular signal and mass transport. Nature Chemistry 1–9 (2024).

[23] Dai, Y., You, L. & Chilkoti, A. Engineering synthetic biomolecular condensates. Nature Reviews Bioengineering 1, 466–480 (2023).

[24] Tran, M. P. et al. A dna segregation module for synthetic cells. Small 19, 2202711 (2023).

[25] Agarwal, S., Klocke, M. A., Pungchai, P. E. & Franco, E. Dynamic self-assembly of compartmentalized dna nanotubes. Nature communications 12, 3557 (2021).

[26] Zhan, P., Jahnke, K., Liu, N. & Göpfrich, K. Functional dna-based cytoskeletons for synthetic cells. Nature Chemistry 14, 958–963 (2022).

[27] Jahnke, K., Huth, V., Mersdorf, U., Liu, N. & Göpfrich, K. Bottom-up assembly of synthetic cells with a dna cytoskeleton. ACS nano 16, 7233–7241 (2022).

[28] Green, L. N., Amodio, A., Subramanian, H. K., Ricci, F. & Franco, E. ph-driven reversible self-assembly of micron-scale dna scaffolds. Nano letters 17, 7283–7288 (2017).

[29] Illig, M. et al. Triggered contraction of self-assembled micron-scale dna nanotube rings. Nature Communications 15, 2307 (2024).

[30] Langecker, M. et al. Synthetic lipid membrane channels formed by designed dna nanostructures. Science 338, 932–936 (2012).

[31] Zhang, Y. et al. Dynamic dna structures. Small 15, 1900228 (2019).

[32] Yu, Z. et al. A self-regulating dna rotaxane linear actuator driven by chemical energy. Journal of the American Chemical Society 143, 13292–13298 (2021).

[33] Centola, M. et al. A rhythmically pulsing leaf-spring dna-origami nanoengine that drives a passive follower. Nature nanotechnology 19, 226–236 (2024).

[34] Andersen, E. S. et al. Self-assembly of a nanoscale dna box with a controllable lid. Nature 459, 73–76 (2009).

[35] Praetorius, F. et al. Biotechnological mass production of dna origami. Nature 552, 84–87 (2017).

[36] Geary, C., Rothemund, P. W. & Andersen, E. S. A single-stranded architecture for cotranscriptional folding of rna nanostructures. Science 345, 799–804 (2014).

[37] Li, M. et al. In vivo production of rna nanostructures via programmed folding of single-stranded rnas. Nature communications 9, 2196 (2018).

[38] Geary, C., Grossi, G., McRae, E. K., Rothemund, P. W. & Andersen, E. S. Rna origami design tools enable cotranscriptional folding of kilobase-sized nanoscaffolds. Nature chemistry 13, 549–558 (2021).

[39] Geary, C. W. & Andersen, E. S. Design principles for single-stranded rna origami structures, Vol. 20, 1–19 (International Society for Nanoscale Science, Computation and Engineering, 2014).

[40] Gaut, N. J. & Adamala, K. P. Reconstituting natural cell elements in synthetic cells. Advanced Biology 5, 2000188 (2021).

[41] Chakraborty, T. & Wegner, S. V. Cell to cell signaling through light in artificial cell communities: glowing predator lures prey. Acs Nano 15, 9434–9444 (2021).

[42] Chakraborty, T., Bartelt, S. M., Steinkühler, J., Dimova, R. & Wegner, S. V. Light controlled cell-to-cell adhesion and chemical communication in minimal synthetic cells. Chemical communications 55, 9448–9451 (2019).

[43] De Franceschi, N., Hoogenberg, B., Katan, A. & Dekker, C. Engineering ssrna tile filaments for (dis) assembly and membrane binding. Nanoscale (2024).

[44] Stewart, J. M., Geary, C. & Franco, E. Design and characterization of rna nanotubes. ACS nano 13, 5214–5221 (2019).

[45] Autour, A., Westhof, E. & Ryckelynck, M. ispinach: a fluorogenic rna aptamer optimized for in vitro applications. Nucleic acids research 44, 2491–2500 (2016).

[46] Ouellet, J. Rna fluorescence with light-up aptamers. Frontiers in chemistry 4, 29 (2016).

[47] Plaskon, D. et al. Step-by-step regulation of productive and abortive transcription initiation by pyrophosphorolysis. Journal of molecular biology 434, 167621 (2022).

[48] Lakshminarayanaiah, N. Transport phenomena in artificial membranes. Chemical reviews 65, 491–565 (1965).

[49] Lanter, F., Erne, D., Ammann, D. & Simon, W. Neutral carrier based ion-selective electrode for intracellular magnesium activity studies. Analytical Chemistry 52, 2400–2402 (1980).

[50] Dolder, N., Müller, P. & von Ballmoos, C. Experimental platform for the functional investigation of membrane proteins in giant unilamellar vesicles. Soft matter 18, 5877–5893 (2022).

[51] Jahnke, K. & Göpfrich, K. Engineering dna-based cytoskeletons for synthetic cells. Interface Focus 13, 20230028 (2023).

[52] Winfree, E., Liu, F., Wenzler, L. A. & Seeman, N. C. Design and self-assembly of two-dimensional dna crystals. Nature 394, 539–544 (1998).

[53] Rothemund, P. W. et al. Design and characterization of programmable dna nanotubes. Journal of the American Chemical Society 126, 16344–16352 (2004)

[54] Šulc, P., Romano, F., Ouldridge, T. E., Doye, J. P. & Louis, A. A. A nucleotide-level coarse-grained model of rna. The Journal of chemical physics 140 (2014).

[55] Poppleton, E. et al. oxdna: coarse-grained simulations of nucleic acids made simple. Journal of Open Source Software 8, 4693 (2023).

[56] Emamyari, S., Mirzaei, M., Mohammadinejad, S., Fazli, D. & Fazli, H. Impact of flexibility on the aggregation of polymeric macromolecules. The European Physical Journal E 46, 66 (2023).

[57] Arulkumaran, N., Singer, M., Howorka, S. & Burns, J. R. Creating complex protocells and prototissues using simple dna building blocks. Nature Communications 14, 1314 (2023).

[58] Bezanilla, M., Gladfelter, A. S., Kovar, D. R. & Lee, W.-L. Cytoskeletal dynamics: a view from the membrane. Journal of Cell Biology 209, 329–337 (2015).

[59] Deamer, D. W. & Fleischaker, G. R. *(Foreword by Joyce Gerald) Origins of life: the central concepts* (Jones and Bartlett, Boston, 1994).

[60] Benner, S. A. Defining life. Astrobiology 10, 1021–1030 (2010).

[61] Saha, R., Verbanic, S. & Chen, I. A. Lipid vesicles chaperone an encapsulated rna aptamer. Nature communications 9, 2313 (2018).

[62] Peter, B., Levrier, A. & Schwille, P. Spatiotemporal propagation of a minimal catalytic rna network in guv protocells by temperature cycling and phase separation. Angewandte Chemie International Edition 62, e202218507 (2023).

[63] Nilsson, M., Antson, D.-O., Barbany, G. & Landegren, U. Rna-templated dna ligation for transcript analysis. Nucleic Acids Research 29, 578–581 (2001).

[64] Horning, D. P. & Joyce, G. F. Amplification of rna by an rna polymerase ribozyme. Proceedings of the National Academy of Sciences 113, 9786–9791 (2016).

[65] Pettersen, E. F. et al. Ucsf chimerax: Structure visualization for researchers, educators, and developers. Protein science 30, 70–82 (2021).

[66] McRae, E. K. et al. Structure, folding and flexibility of co-transcriptional rna origami. Nature Nanotechnology 1–10 (2023).

[67] Snodin, B. E. et al. Introducing improved structural properties and salt dependence into a coarse-grained model of dna. The Journal of chemical physics 142 (2015).

[68] Ouldridge, T. E., Louis, A. A. & Doye, J. P. Structural, mechanical, and thermodynamic properties of a coarse-grained dna model. The Journal of chemical physics 134 (2011).

[69] Rovigatti, L., Šulc, P., Reguly, I. Z. & Romano, F. A comparison between parallelization approaches in molecular dynamics simulations on gpus. Journal of computational chemistry 36, 1–8 (2015).

[70] Suma, A. et al. Tacoxdna: A user-friendly web server for simulations of complex dna structures, from single strands to origami. Journal of computational chemistry 40, 2586–2595 (2019).

[71] Doye, J. P. et al. in The oxdna coarse-grained model as a tool to simulate dna origami 93–112 (Springer, 2023).

[72] Bohlin, J. et al. Design and simulation of dna, rna and hybrid protein–nucleic acid nanostructures with oxview. Nature protocols 17, 1762–1788 (2022).

[73] Poppleton, E. et al. Design, optimization and analysis of large dna and rna nanostructures through interactive visualization, editing and molecular simulation. Nucleic acids research 48, e72–e72 (2020).

[74] Snodin, B. E. et al. Direct simulation of the self-assembly of a small dna origami. ACS nano 10, 1724–1737 (2016).

[75] Russo, J., Tartaglia, P. & Sciortino, F. Reversible gels of patchy particles: role of the valence. The Journal of chemical physics 131 (2009).

[76] Něcas, D. & Klapetek, P. Gwyddion: an open-source software for SPM data analysis. Central European Journal of Physics 10, 181–188 (2012).

[77] Steger, C. An unbiased detector of curvilinear structures. IEEE Transactions on Pattern Analysis and Machine Intelligence 20, 113–125 (1998). URL 10.1109/34.659930.

[78] Wagner, T., Hiner, M. & Xraynaud. thorstenwagner/ij-ridgedetection: Ridge detection 1.4.0 (2017). URL https://zenodo.org/record/845874.

